# Proliferating CLL cells express high levels of CXCR4 and CD5

**DOI:** 10.1101/2023.12.27.573410

**Authors:** Daniel Friedman, Drshika P Mehtani, Jennifer B Vidler, Piers EM Patten, Robbert Hoogeboom

## Abstract

Chronic Lymphocytic Leukaemia (CLL) is an incurable progressive malignancy of CD5+ B cells with a birth rate between 0.1-1% of the entire clone per day. However, the phenotype and functional characteristics of proliferating CLL cells remain incompletely understood. Here, we stained peripheral blood CLL cells for ki67 and DNA content and found that CLL cells in G1-phase have a CXCR4^lo^CD5^hi^ phenotype, whilst CLL cells in S/G2/M-phase express high levels of both CXCR4 and CD5. Induction of proliferation *in vitro* using CD40L stimulation results in high ki67 levels in CXCR4^lo^CD5^hi^ cells with CXCR4 expression increasing as CLL cells progress through S and G2/M-phases, whilst CXCR4^hi^CD5^lo^ CLL cells remained quiescent. Dye dilution experiments revealed an accumulation of Ki67^hi^ divided cells in the CXCR4^hi^CD5^hi^ fraction. In Eμ-TCL1 transgenic mice, the CXCR4^hi^CD5^hi^ fraction expressed high levels of ki67 and was expanded in enlarged spleens of diseased animals. Human peripheral blood CXCR4^hi^CD5^hi^ CLL cells express high levels of IgM and the chemokine receptors CCR7 and CXCR5 and migrated most efficiently towards CCL21. We found higher levels of CXCR4 in patients with progressive disease and the CXCR4^hi^CD5^hi^ fraction was expanded upon clinical relapse. Thus, this study defines the phenotype and functional characteristics of proliferating CLL cells identifying a novel subclonal population that underlies CLL pathogenesis and may drive clinical outcome.

## Introduction

Chronic lymphocytic leukaemia (CLL) is characterised by an expansion of malignant CD5+ B cells in the peripheral blood (PB), bone marrow (BM) and secondary lymphoid tissues. CLL cells proliferate substantially with between 0.1-1% of the clone renewed per day^1^. In the clinic, CLL patients display a heterogeneous course. CLL cases expressing B cell receptors (BCRs) with unmutated *IGHV* (U-CLL) have a worse prognosis and more frequently require treatment, whilst CLL with mutated *IGHV* (M-CLL) genes may have stable disease for years, reflecting higher net CLL cell birthrates in U-CLL patients^2,3^.

*In vivo* deuterium labelling revealed that proliferation rates of CLL cells are highest in the lymph nodes (LNs), where CLL cells display more prominent BCR activation and resident immune and stromal cells present key co-stimulatory ligands such as antigen and CD40 ligand (CD40L)^4,5^. Following stimulation and proliferation in LNs, CLL cells may emigrate to the PB, where cells may acquire a quiescent state until either tissue re-entry and re-stimulation or death ensues. Whilst the PB compartment is largely deemed quiescent, the discovery of subpopulations of newly-born and quiescent cells in the PB indicates a spectrum of cell states among circulating CLL cells in the periphery^6^. At one end are activated recent LN emigrants, expressing low levels of the chemokine C-X-C motif receptor 4 (CXCR4) and high levels of CD5, often used as a proxy for proliferating cells in LNs^7–9^. At the other end, the fraction of cells with high CXCR4 and low CD5 is enriched in quiescent cells. However, a recent study integrating bulk and single-cell RNA sequencing of LN and PB CLL cells identified three major cell states in both compartments: quiescent, activated, and proliferating^10^, suggesting the current CXCR4/CD5 model may not capture all cell states.

Identifying which cells proliferate is important for developing therapeutic strategies that target the most relevant cells and we therefore set out to identify and functionally characterise actively dividing cells from the PB of CLL patients. We find that PB CLL cells in S, G2 and M phases uniformly express high levels of CXCR4 and CD5. These CXCR4^hi^CD5^hi^ cells are enriched for highly proliferative cells upon stimulation *in vitro* and have elevated levels of chemokine receptors, facilitating efficient migration in response to LN homing signals. Finally, we observed increased levels of CXCR4 and CD5 in PB CLL samples at relapse. Together, this study provides novel insights into the nature of proliferating cells in CLL as well as new starting points to develop biomarkers for disease progression and response to therapies that target proliferative signals.

## Materials and Methods

### Patients and samples

This research was supported by material from the King’s College Denmark Hill Haematology Biobank (18/NE/0141). Cryopreserved PB samples were obtained from CLL patients with written informed consent in accordance with the Declaration of Helsinki. PB and LN fine needle aspirates were taken as previously described^11^.

### Mice

Animal experiments were approved by the UK Home Office. BM, spleen and PB were harvested from aged Eμ-*TCL1* mice (>10 months). Mice harbouring <50% CD19+CD5+ splenic lymphocytes were excluded from the study. Single cell suspensions from tissue were obtained by mechanical disruption and passing cells through 40μm cell strainers (Corning). Red blood cells were lysed in blood and spleen samples using RBC-lysis buffer (Biolegend).

### In vitro proliferation assays

Primary CLL cells were cultured for up to 9 days in Iscove’s Modified Dulbecco’s Medium (IMDM; Gibco) supplemented with 10% Fetal bovine serum (Gibco), 1% Penicillin/Streptomycin and 1% L-glutamine (both Sigma). Where indicated, CLL PBMCs were cocultured on irradiated NIH3T3 fibroblasts stably transfected with CD40 ligand or parental cells with or without interleukin-4 (IL-4, 10ng/ml, Peprotech) and IL-21 (25ng/ml, Peprotech). To track proliferation histories, PBMCs (10x10^6^/ml) were labelled with 0.5μM CMFDA (ThermoFisher) in PBS for 20 minutes at 37°C prior to seeding.

### Flow cytometry

Human PBMCs were treated with Fc blocking solution (Human TruStain FcX, Biolegend) and stained with the following fluorescent antibodies from Biolegend (unless otherwise stated): anti-CD19 (HIB19), anti-CD5 (L17F12), anti-CXCR4 (12G5), anti-CCR7 (G043HD), anti-CXCR5 (J252D4), anti-IgM (MHM-88), anti-CD49d (9F10), anti-AID (BD Biosciences, EK2-5G9), anti-Ki67 (11F6) and anti-NFAT2 (7A6). Mouse primary cells were treated with anti-mouse CD16/32 (Biolegend) and stained with fluorescent antibodies: anti-CD19 (6D5), anti-CD5 (53-7.3), anti-CXCR4 (L276F12) and anti-Ki67 (11F6). A fixable viability dye (eBioscience) was used to exclude dead cells. Cells were fixed with 4% PFA and washed with PBS prior to acquisition. For intracellular staining, cells were fixed and permeabilised using the FoxP3/Transcription factor staining kit (eBioscience) and stained overnight for Ki67 at 4°C. To discriminate between cell cycle stages, permeablised cells were treated with 100μg/ml RNase A (Sigma) for 15 mins at 37°C followed by labelling with 0.5μg/ml DAPI (Sigma) for 10 mins at room temperature. Cells were acquired on a BD LSR Fortessa and data analysed using Flowjo software (TreeStar, V10). For imaging flow cytometry, cells were stained as above and analysed on an ImagestreamX MK-II (Amnis). Brightfield and fluorescent images were captured using 40× zoom, and data were analysed using IDEAS (Merck). Cell debris and doublets were excluded prior to analysis.

Ki67^lo^DAPI^hi^ populations were excluded from the S/G2/M gate to account for the presence of shadow doublets^12^.

### Migration assays

Migration assays were performed using 5-μm pore polycarbonate transwell inserts in 24-well plates (Corning). Transwell filters were coated overnight with 2.5μg/ml ICAM-1 (R&D systems) in PBS and blocked with 2% BSA/PBS for 2hrs. 0.5% BSA plus 0.1/1 μg/mL CCL21 (R&D systems) was added to basolateral chambers, and 0.5x10^6^ PBMCs were transferred into the apical chambers and incubated for 2 hours at 37°C/5% carbon dioxide. Time-matched controls were incubated in ICAM-1 coated wells in parallel with the migration assays. After 2hrs, 40mM EDTA/PBS was added to each well/filter followed by staining with fluorescent antibodies against CD19, CD5 and CXCR4 as described above.

### Statistical analysis

Statistical analyses were carried out using GraphPad Prism (version 10). Shapiro–Wilk normality tests were used to evaluate the distribution of values. Data are presented as means±standard deviation.

## Results

### Proliferating CLL cells in the blood express high levels of CXCR4 and CD5

The low frequency of proliferating cells (∼0.1-1% of the leukeamic clone) has made it challenging to identify and characterise actively dividing cells in the periphery. Here, we stained PB samples from U-CLL and M-CLL patients for DNA content and Ki67 expression to distinguish cells in G0 (2n DNA, ki67^lo^), G1 (2n DNA, ki67^hi^) and S/G2/M (4n DNA, Ki67^hi^) phases (Fig.1a). Consistently, we detected a fraction of cells in S/G2/M phases (0.24±0.17%) in all 20 patients analysed (Fig.1b).

**Figure 1.**
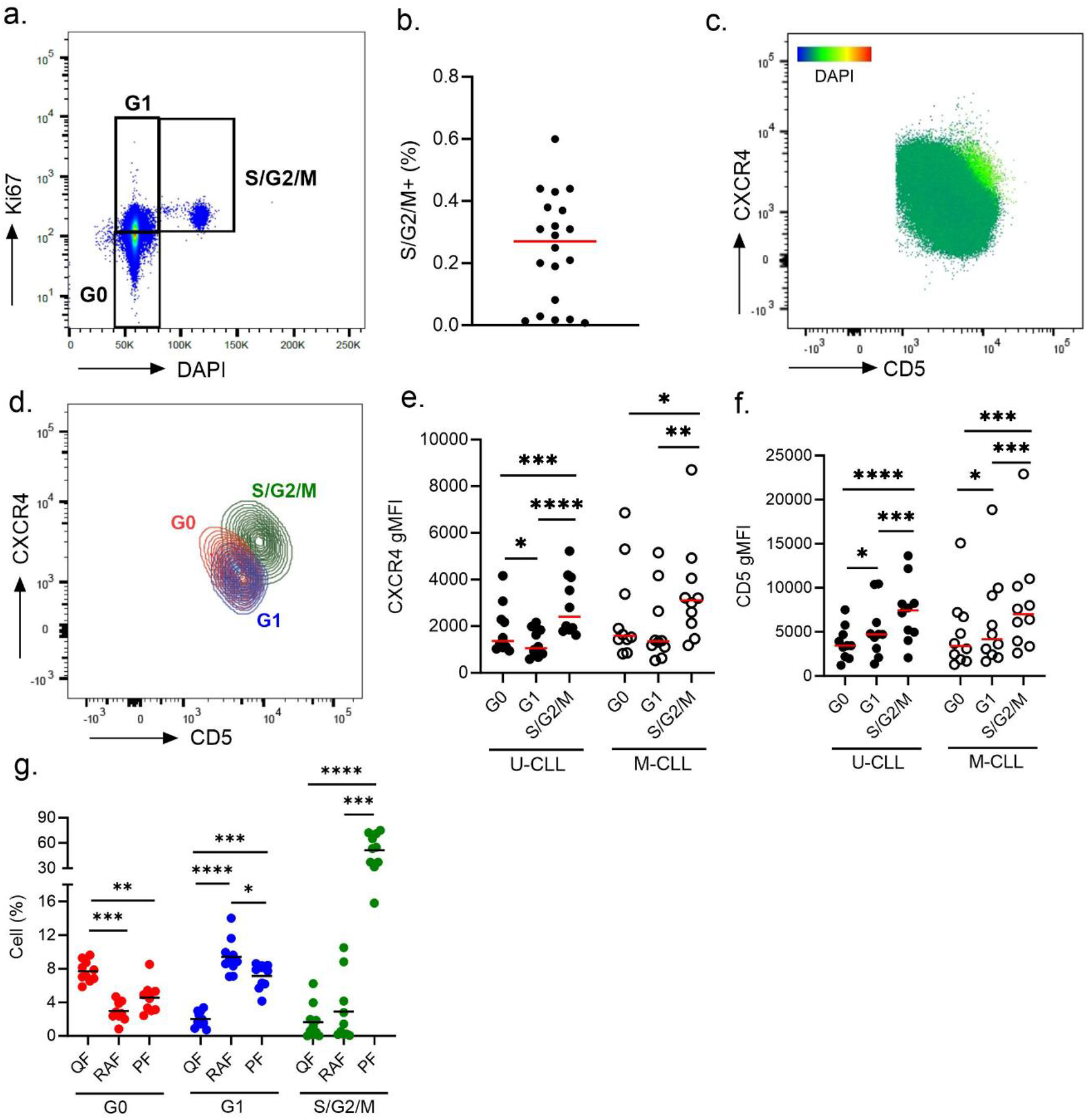
Proliferating CLL cells in the blood express high levels of CXCR4 and CD5. (a) Representative scatter plot of Ki67 and DAPI staining on dividing CLL cells highlighting cell fractions in G0, G1 and S/G2/M phases. Quantification of the percentage of cells in S/G2/M phases. n=20 (c) CXCR4 vs CD5 scatter plot overlaid with cells in S/G2/M-phase which can be observed localised to the CXCR4^hi^CD5^hi^ fraction. DAPI fluorescence intensities are displayed as a heat map scale. (d) Representative contour plots showing CXCR4 and CD5 expression on G0, G1 and S/G2/M fractions as gated in (a). Quantification of (e) CXCR4 and (f) CD5 gMFIs on cells in G0, G1 and S/G2/M phases for both M- and U-CLL patients. Each data point represents a single patient, n=10. (g) The percentage of cells in G0, G1 and S/G2/M in different cell fractions were quantified. Each data point represents a single patient, n=20. gMFI, geometric mean fluorescence intensities. Statistical significance of data was calculated using a repeated measures (RM) one-way ANOVA with Tukey’s multiple comparisons. *p <0.05, **p <0.01, ***p <0.001, ****p <0.0001. QF: Quiescent Fraction, RAF: Recent Activated Fraction, PF: Proliferating Fraction.

*In vivo* deuterium labelling studies have identified that the CXCR4^lo^CD5^hi^ cell fraction is enriched in recently proliferated cells, whilst CXCR4^hi^CD5^lo^ phenotypes represent older quiescent CLL cells. To investigate which cell fraction contains actively dividing cells, we stained PB CLL cells for CXCR4, CD5 and DNA content. Surprisingly, cells with >2n DNA map to a fraction with high CXCR4 and high CD5 (Fig.1c), indicating that actively dividing CLL cells have a distinct surface phenotype. To demonstrate that DAPI^hi^ cells are indeed in the cell cycle, we visualised the nuclei of CXCR4^hi^CD5^hi^ cells using imaging flow cytometry and observed cells in mitosis (Supp fig.1a).

To interrogate the phenotypes of quiescent and proliferating CLL cells further, we overlayed the G0, G1 and S/G2/M fractions revealing distinct CXCR4 and CD5 expression profiles (Fig.1d) with lowest levels of CXCR4 on cells in G1 in U-CLL and highest CXCR4 expression levels on cells in S/G2/M for both U-CLL and M-CLL (Fig.1e). CD5 expression increased as cells progressed through the cell cycle peaking in S/G2/M phases for both M-CLL and U-CLL cells (Fig.1f).

To quantify the percentage of G0, G1 and S/G2/M cells within different cell fractions based on CXCR4 and CD5 densities, gates comprising 5% of the PB bulk population were drawn to examine previously characterised CXCR4^hi^CD5^lo^ (quiescent fraction, QF) and CXCR4^lo^CD5^hi^ (recent activated fraction, RAF)^6^ and the CXCR4^hi^CD5^hi^ subpopulation that harbours cells in S/G2/M phase (proliferating fraction, PF) (Supp fig.1b). The highest percentage of cells in G0 were residing in the QF whilst the RAF contained the highest percentage of cells in G1 (Fig.1g). Strikingly, most cells in S/G2/M phases were residing in the PF. Altogether, these data demonstrate that the RAF is enriched for G1 cells whilst mitotic CLL cells express high levels of both CXCR4 and CD5.

### CXCR4^hi^CD5^hi^ cells are highly proliferative in the TCL1 mouse model for CLL

PB cell fractions are thought to represent a shadow of signalling events in tissue where CLL cell activation and proliferation predominantly occur^4,5^. Thus, we next compared proliferating cells in PB and tissues from aged transgenic *Eμ-TCL1* mice. We found high levels of Ki67 on CD19^+^CD5^+^ murine leukemia cells from the PB and spleen whilst leukemic cells in the BM expressed significantly lower levels, suggesting that in this mouse model proliferating cells are not sequestered in the BM compartment (Supp fig.2a). Co-staining PB and spleen cells with ki67 and DAPI revealed similar G1 and S/G2/M fraction sizes in both compartments (Supp fig.2b-d). Of note, the frequency of cells in S/G2/M is higher than observed in humans, confirming the aggressive nature of the *Eμ-TCL1* mouse model. To examine CXCR4 and CD5 expression on cells in different stages of the cell cycle, we overlayed the G0, G1 and S/G2/M fractions revealing distinct CXCR4 and CD5 expression profiles for these subpopulations (Fig.2a). S/G2/M cells expressed the highest levels of CXCR4 and CD5 (Fig.2b and c) mirroring our findings in human CLL cells.

**Figure 2.**
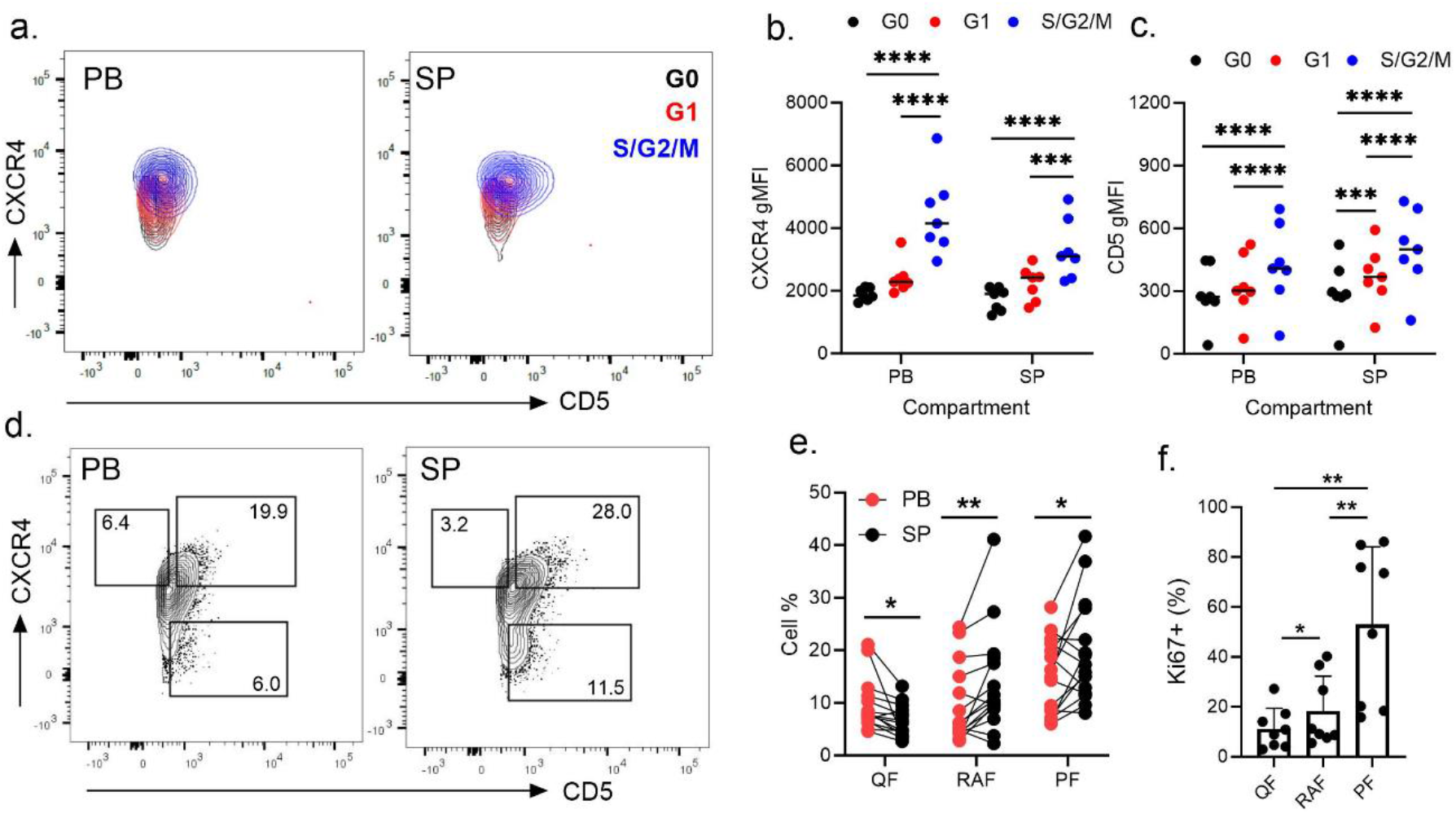
CXCR4^hi^CD5^hi^ cells are highly proliferative in the Eμ-TCL1 mouse model for CLL. (a) Representative contour plots showing CXCR4/CD5 expression on G0 (black), G1 (red) and S/G2/M (blue) fractions as gated in supp. Fig 2b. Quantification of (b) CXCR4 and (c) CD5 gMFIs on cells in G0, G1 and S/G2/M phases from the peripheral blood (PB) and spleen (SP) compartments. Each data point represents a single mouse, n=7. (d) Representative contour plots comparing CXCR4/CD5 profiles in cells from the PB and spleen. Gates drawn reflect the Quiescent fraction (QF, CXCR4^hi^CD5^lo^), Proliferating Fraction (PF, CXCR4^hi^CD5^hi^) and Recent Activated Fraction (RAF, CXCR4^lo^CD5^hi^) with the cell percentage values shown. Gates were drawn on the PB and extrapolated onto spleen cell plots. (e) Quantification of cell percentages in the three different fractions as gated in (d). Data shows matched values, where each data point represent an individual mouse, n=15. (f) Quantification of Ki67+ cells in each cell fraction in the spleen. Each data point represents a single mouse, n=8. Data shows mean±SEM. gMFI, geometric mean fluorescence intensities. Statistical significance of data were calculated using a RM one-way ANOVA with Tukey’s multiple comparisons or paired parametric t-tests *p <0.05, **p <0.01, ***p <0.001, ****p <0.0001.

Leukemic *Eμ-TCL1* mice present with enlarged spleens. To investigate if spleen enlargement is associated with accumulation of cells with distinct phenotypes, we compared CXCR4 and CD5 profiles of spleen and PB CLL cells and observed expansions of both the RAF and the PF in the spleen, with the latter significantly enriched with Ki67+ cells (Fig.2d-f). Unique to the BM compartment we detected an expansion of the QF, in line with this compartment harbouring more quiescent cells (Supp fig.2e). Altogether, these data suggests that proliferating cells in the *Eμ*-*TCL1* mouse model have a CXCR4^hi^CD5^hi^ phenotype accumulating predominantly in the spleen.

*Eμ-TCL1* mice exhibit similar features to aggressive human CLL and the expansion of a Ki67^hi^CXCR4^hi^CD5^hi^ fraction in the spleen may reflect enhanced proliferation rates. Consistent with this idea, examination of human LN cells from a patient with aggressive CLL (de novo TP53-mutated U-CLL patient, requiring immediate treatment) revealed a striking expansion of a Ki67^hi^CXCR4^hi^CD5^hi^ fraction in the LN compared to the PB compartment (Supp fig.3a and b). Conversely, we observed a massively expanded RAF in the PB whilst only a negligible RAF was detected in the LN, suggesting the RAF in PB may emanate from CXCR4^hi^ populations in tissue. Collectively, these findings support the hypothesis that an expanded CXCR4^hi^CD5^hi^ fraction in tissue may reflect enhanced cell proliferation rates.

### Modulation of CXCR4 expression in proliferating cells

Our observation that G1 U-CLL cells express low levels of CXCR4 whilst S/G2/M cells have CXCR4^hi^CD5^hi^ phenotypes, suggests that the surface phenotype of CLL cells may change when cells proliferate. To investigate how CXCR4 and CD5 expression varies on cells in different phases of the cell cycle, robust proliferation of human PB U-CLL cells was induced using CD40-ligand (CD40L) expressing fibroblasts alone or in the presence of interleukin (IL)-4 and IL-21. After 48hrs on CD40L alone, when bulk ki67 levels are still low, U-CLL cells displayed significantly reduced surface CXCR4 expression, whilst CD5 levels remained unchanged. Addition of IL-4 and IL-21 resulted in even lower CXCR4 surface levels compared to CD40L alone. Contrastingly, in unstimulated cells, CXCR4 levels increased three-fold whilst CD5 surface levels increased 1.3-fold (Supp fig.4a-c). We observed no significant fluctuations in CXCR4 and CD5 expression over the initial 48hrs of stimulation in cultures of M-CLL cells. Staining with DAPI and Ki67 revealed that after 48hrs the G1 fraction expressed the lowest levels of CXCR4, with CD5 levels increasing and peaking on S/G2/M cells (Fig.3a).

**Figure 3.**
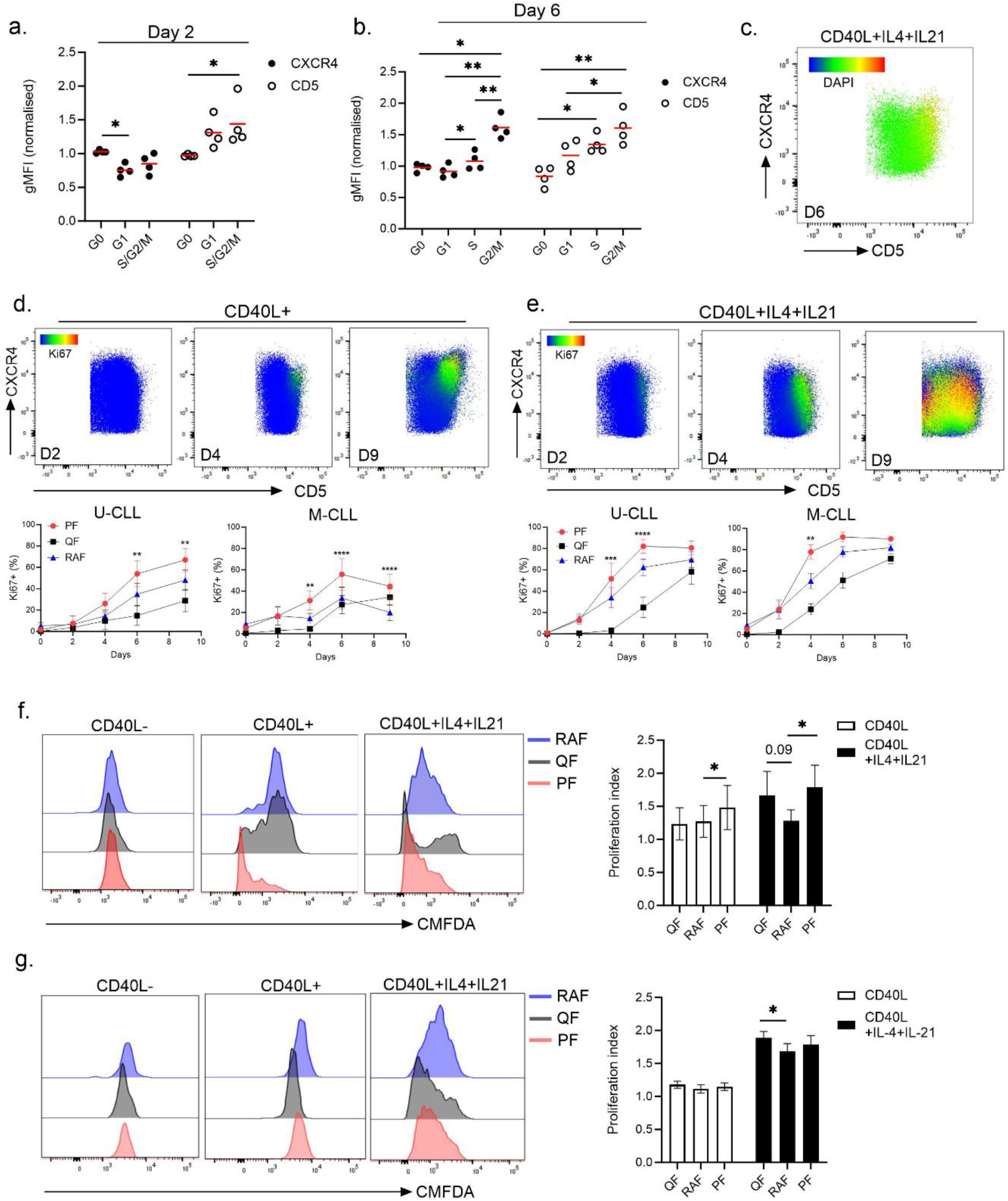
CXCR4 and CD5 are upregulated when CLL cells progress through the cell cycle. (a) Quantification of CXCR4 and CD5 expression levels of U-CLL cells in G0, G1 and S/G2/M phases when stimulated on CD40L-expressing fibroblasts with interleukin (IL) 4 and IL-21 after 2 and (b) 6 days. (n=4) (c) Representative CXCR4/CD5 scatter plot overlaid with DAPI gMFIs where DAPI^hi^ cells can be observed localised to the CXCR4^hi^CD5^hi^ fraction. DAPI fluorescence intensities are displayed as a heat map scale. Human U-CLL and M-CLL cells were seeded on CD40L-expressing fibroblasts alone (d) or in the presence of IL-4 and IL-21 (e) for 9 days and samples assessed for Ki67 expression on days 0, 2, 4, 6 and 9. Representative scatter plots from an U-CLL patient of CXCR4 and CD5 profiles from days 2, 4 and 9 are shown with Ki67 gMFIs overlaid as a heatmap statistic. Quantification of Ki67 expression in M-CLL and U-CLL cells in different cell fractions over 9 days are shown in lower panels (n=8). To track cell proliferation histories within fractions, (f) U- and (g) M-CLL primary CLL cells were stained with CMFDA prior to cell stimulation. Representative histogram plots from cells on day 9 are shown (left panels) with quantified proliferation indices for both U-CLL and M-CLL patients (right panel; n=8). Data shows mean±SEM. gMFI, geometric mean fluorescence intensities. Statistical significance of data were calculated using a RM one-way ANOVA with Tukey’s multiple comparisons *p <0.05, **p <0.01, ***p <0.001, ****p <0.0001. QF: Quiescent Fraction, RAF: Recent Activated Fraction, PF: Proliferating Fraction.

After 6 days, we observed a stepwise increase in both CXCR4 and CD5 expression as cells progressed from G1 into S and G2/M phases (Fig.3b) and the DAPI^hi^ subpopulation mapped to the PF (Fig.3c), mirroring our findings in freshly thawed PBMCs. To investigate if the three cell states observed in PB can be recapitulated *in vitro*, we gated the QF, RAF and PF as previously described. From 4 days onwards of CD40 activation, when CLL cells are proliferating, the RAF and the PF were significantly enriched with Ki67^+^ cells in U-CLL cultures, while ki67 expression remained low in the QF (Fig.3d). Proliferation was less robust in cultures with M-CLL cells, showing significant enrichment of ki67 in the PF, but low ki67 in the RAF and the QF. Combining CD40L with IL-4 and IL-21, increased the frequency of Ki67+ cells observed by 48hrs in the RAF and PF and by day 6 the majority of cells in these fractions were Ki67+ in both U-CLL and M-CLL (Fig.3e). Of note, the QF contained the lowest percentage of Ki67+ cells. However, Ki67+ cells in the QF increased by day 6 of culture, matching the levels of the RAF and PF by day 9 (Fig.3e).

To track cell proliferation histories over 9 days in culture, cells were stained with the fluorescent dye CMFDA prior to seeding. The PF was significantly enriched with proliferated cells in U-CLL cultures in response to either CD40 stimulation alone or in combination with cytokines. Additionally, we observed an increase in divided cells in the QF compared to the RAF when cells were stimulated with both CD40L and cytokines, although this effect did not reach statistical significance (*p*=0.09; fig.3f). In contrast, cell proliferation was more subdued in M-CLL cells in response to CD40L alone and when combined with cytokines. Whilst we observed increased cell proliferation, proliferating cells accumulated in the QF with no significant increase detected in the PF (Fig.3g). Thus, our data demonstrates that whilst activating stimuli initially lead to a strong downregulation of CXCR4 reflecting cell entry into G1, it quickly resurfaces when cells enter S/G2/M phases, revealing an intricate relationship between activation, proliferation and migration potential.

### Proliferating CLL cells have a pro-migratory phenotype

Whilst the RAF and QF have been well characterised little is known regarding the phenotype and fate of PB CXCR4^hi^CD5^hi^ cells. *In vivo* deuterium labelling revealed a correlation between proliferation and high expression of surface IgM^13^, important for CLL cell re-stimulation with antigen. We found IgM was expressed at high levels on both the PF and RAF in U-CLL patients with lowest IgM levels on the QF (Fig.4a). In M-CLL, bulk IgM levels were lower, as expected^14^, with no significant differences between fractions. Expression of activation-induced cytidine deaminase (AID) is associated with ki67 expression in tissue^15^. To investigate what fraction(s) express(es) AID we analysed intracellular AID levels using flow cytometry. AID expression was highest in the RAF in both U- and M-CLL patients although U-CLL expressed higher levels of AID in the PF fraction compared to the QF (Fig.4b).

**Figure 4.**
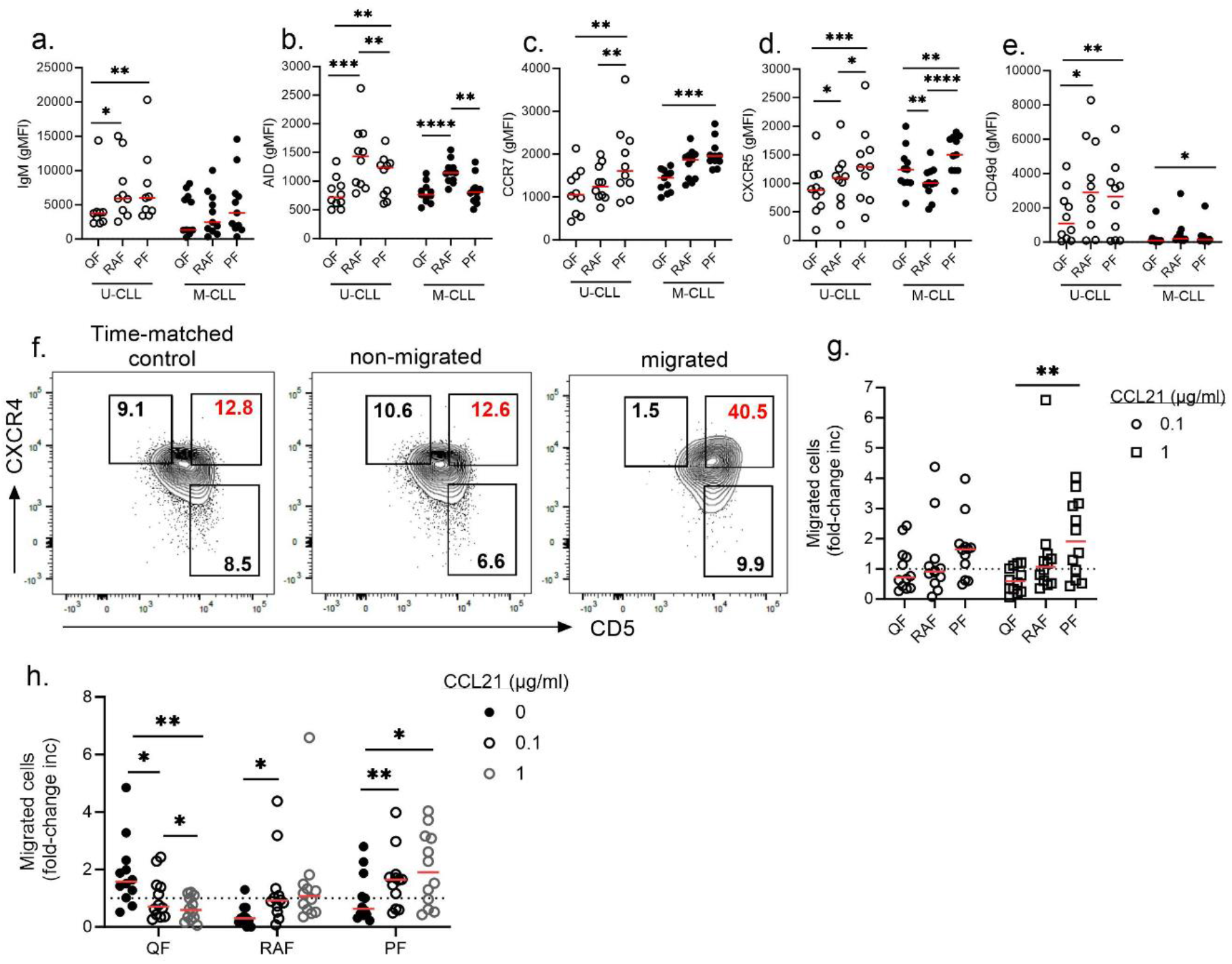
Proliferating cells have a pro-migratory phenotype. Expression levels of (a) surface IgM (b) AID CCR7 (d) CXCR5 and (e) CD49d were quantified on U- and M-CLL cells. (f) PBMCs from U-CLL patients were placed in transwell migration chambers and incubated in the absence or presence of increasing concentrations of CCL21. After 2hrs cells were harvested from the top chamber (non-migrated) and bottom chamber (migrated) and cells were stained with CD19, CD5, CXCR4 and CD5. CXCR4 and CD5 gates were drawn on time matched controls and extrapolated on migrated and non-migrated fractions to quantify changing fraction sizes. Representative scatter plots are shown from a U-CLL patient. Fraction percentages are shown within each gate with the changing CXCR4^hi^CD5^hi^ values shown in red. (g) Quantification of the fold-change increase in fraction size in U-CLL patients of migrated cells at both lower (0.1μg/ml) and higher (1μg/ml) CCL21 concentrations (n=12). Migrated cells = number of migrated cells with chemokine/number of migrated cells in the absence of chemokine. (h) Comparison of the fold-change increase in fraction size with increasing CCL21 concentrations (n=12). Data points represent individual patients. Statistical significance of data were calculated using a RM one-way ANOVA with Tukey’s multiple comparisons *p <0.05, **p <0.01, ***p <0.001, ****p <0.0001. QF: Quiescent Fraction, RAF: Recent Activated Fraction, PF: Proliferating Fraction.

Previous analysis identified a fraction of CLL cells expressing high levels of IgM and CXCR4^16^, which may represent a dangerous cell fraction in the PB primed for homing to tissue. Thus, we next assessed expression of receptors important for migration into and positioning within LNs. Strikingly, the PF showed the highest expression of both CCR7 and CXCR5 in both U-CLL and M-CLL patients across all fractions assessed (Fig.4c and d) although no difference was observed in bulk levels between U-CLL and M-CLL (Supp Fig.5a and b). CD49d was expressed at similar levels on both the RAF and PF with lowest levels observed in the QF (Fig.4e). Thus, CXCR4^hi^CD5^hi^ cells express high levels of both chemokine and adhesion receptors.

CCR7 has been strongly implicated in LN homing^17^ and we next assessed whether higher levels of CCR7 on CXCR4^hi^CD5^hi^ cells confer a greater propensity to migrate towards CCL21. When examining the bulk population, U-CLL cells migrated more efficiently in transwell assays than M-CLL cells at higher concentrations of CCL21, whilst no differences were detected at lower CCL21 concentrations (Supp Fig.5c). To assess which cell fraction migrated most efficiently towards CCL21, gates were set on time-matched controls to account for fluctuations in CXCR4 expression levels. Importantly, both CXCR4 and CD5 expression levels did not fluctuate in response to cells binding to either CCL21 or ICAM-1 (Supp fig.5d-g). Migrated cells were significantly enriched with cells from the PF compared to RAF and QFs in U-CLL samples (2.0±1.2, 1.4±1.6 and 0.6±0.4 respectively; fig.4f and g). Moreover, whilst RAF cells migrated in response to CCL21 we uniquely observed a dose dependent migratory effect of CCL21 on the PF whilst conversely, cells with a QF phenotype significantly declined with increasing concentrations of CCL21 (Fig.4h). Contrastingly, in M-CLL samples, the RAF showed the largest expansion, while a simultaneous increase in cells from the PF failed to reach statistical significance (*p=*0.08; supp fig.5h). Altogether these data highlight the PF in PB with heightened migratory potential towards CCL21, expressing high levels of IgM, CD49d, CCR7, CXCR5 and CXCR4, facilitating LN homing and re-stimulation.

### Disease progression is associated with increased CXCR4 and CD5 levels

To investigate whether the presence of proliferating cells in the periphery is indicative of proliferation rates in tissue, we next assessed whether patients with progressing disease show larger proliferating cell fractions in the PB compared to more indolent patients. Surprisingly, no differences were detected in the size of the S/G2/M fractions between an indolent and a progressing cohort (*p=*0.361), suggesting that irrespective of disease state the fraction of dividing cells in the blood remains constant (Fig.5a). Consistent with CXCR4 being a negative prognostic marker in CLL, we observed higher expression levels of surface CXCR4 on the bulk population of progressing patients and a trend for higher levels of CD5 (*p*=0.07; Fig.5b). S/G2/M cells in progressing patients expressed higher levels of CXCR4 compared to S/G2/M cells in indolent patients with a strong trend for increased expression of CD5 (*p=*0.056; Fig.5c).

**Figure 5.**
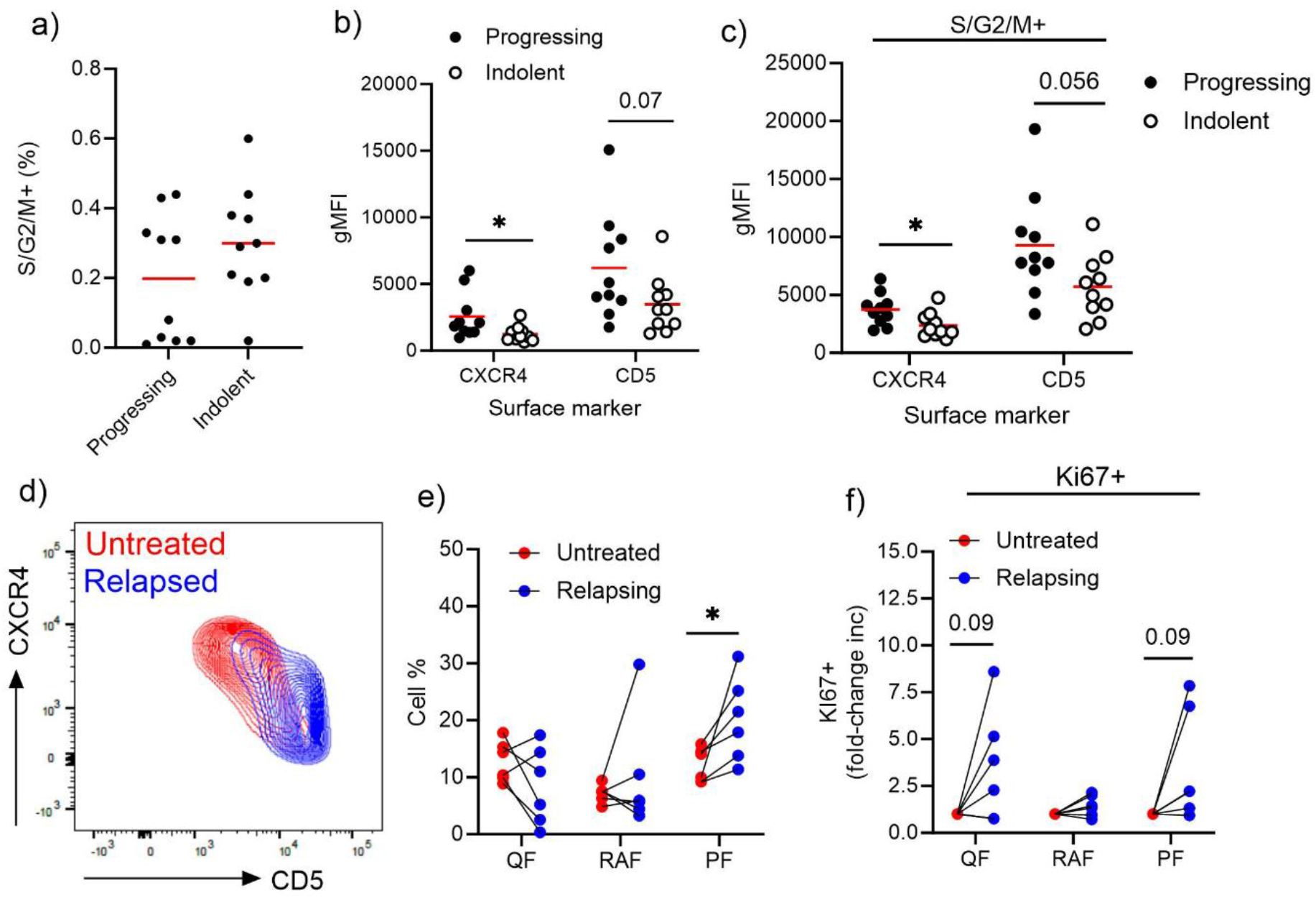
Disease progression is associated with increased CXCR4 and CD5 levels. (a) PB CLL cells from progressing and indolent untreated cohorts were stained with antibodies against Ki67 and DAPI to quantify S/G2/M fractions. CXCR4 and CD5 levels on PB CLL cells from both progressing and indolent patients were quantified on the (b) bulk population and on the (c) S/G2/M fraction. Data points represent individual patients. CXCR4 and CD5 expression profiles were compared in CLL patients before treatment (red) and at relapse (blue). Representative example plots of a single patient is shown. (e) Quantification of cell fraction percentages in 6 patients prior to treatment (red) and in matched relapsing samples (blue). (f) Quantification of the fold-change increase in the percentage of Ki67+ cells in cell fractions in untreated and matched relapsing samples. Statistical significance of data were calculated using unpaired or paired parametric t-tests where appropriate. *p <0.05. QF: Quiescent Fraction, RAF: Recent Activated Fraction, PF: Proliferating Fraction.

We next assessed whether changes in the PF occur within patients progressing over time rather than measuring at a single timepoint. We compared fraction sizes in PB CLL samples from patients relapsing on treatment to samples taken prior to treatment and observed expansion of the PF in all six patients when relapsing (Fig.5d and e), with a trend for increased Ki67 expression in both the PF and QFs at relapse (*p*=0.09) (Fig.5f). Altogether, we conclude that higher CXCR4 and CD5 levels reflect higher net cell birth rates and disease progression.

## Discussion

Deuterium incorporation studies highlighted both quiescent and newly born recent LN emigrants in the PB however identifying actively dividing cells has proven challenging. Here, we combined DNA labelling with Ki67 staining to distinguish cells in G0, G1 and S/G2/M phases and found a small fraction of CLL cells in S/G2/M phase readily detectable in the PB of all patients, regardless of *IGHV* mutation status or disease course. On average, 0.24±0.17% of CLL cells were in S/G2/M phase, a value surprisingly consistent with previous estimations for daily tumour proliferation rates of 0.1-1%^1^ and 0.2%^13^ and not much lower than recently reported for LN CLL cells (0.4-1%)^10^. As CLL patients have ≥5*10^9^ CLL cells/L in the periphery, proliferation of 0.2% PB CLL cells may represent a substantial part of cell turnover.

Recent single-cell transcriptomic analysis of both LN and PB CLL cells distinguished three major cell states, quiescent, activated and proliferating^10^, whilst the CXCR4/CD5 model describes recently divided and quiescent cell fractions^6^. Our data demonstrate that actively proliferating CLL cells uniformly express high levels of both CXCR4 and CD5, indicating the CXCR4^hi^CD5^hi^ fraction contains actively proliferating cells, whilst the RAF fraction contains few cells in S/G2/M-phase. Likely, the higher enrichment of ^2^H-labelled DNA in the RAF reflects proliferation histories rather than active proliferation. Of note, the CXCR4^hi^CD5^hi^ fraction contained ^2^H-labelled cells *in vivo*^18^ in agreement with our observation of mitotic cells in this fraction. Separately, *in vivo* pulse-chase labelling with deuterated glucose revealed a correlation between proliferation and high IgM levels^13^, consistent with our finding of high IgM levels in the PF. Previous studies labelled the RAF as a proliferating fraction based on high Ki67 expression^5,6^. However, the ability of Ki67 to discriminate actively dividing cells is debated^19,20^. Whilst absent in G0 cells (Gerde 1984), Ki67 mRNA and protein levels increase in G1 cells that have not (yet) committed to the cell cycle^21,22^. Based on our data we conclude that the RAF is enriched in G1 cells. Indeed, proteins regulating the G1-S transition, such as *Cyclin E1, Cyclin D2* and MCM6^23–25^ are upregulated in the RAF^6^. Reconciling these studies and our data, we propose that CXCR4^hi^CD5^hi^, CXCR4^lo^CD5^hi^ and CXCR4^hi^CD5^lo^ PB cell fractions may represent proliferating, activated and quiescent fractions respectively.

Among PB CLL cells, only a minor subfraction of activated cells in G1 may progress to the proliferating state where the life cycle of a single CLL cell starts over, whilst the majority may exit to G0. This is in line with recent RNA velocity analysis of single cell transcriptomics data indicating LN CLL cells transverse unidirectionally from proliferating to an activated state before entering a final quiescent state^10^. To examine fates after cell division, we tracked proliferation histories *in vitro*, revealing that the most divided cells by day 9 contained both quiescent and proliferating phenotypes, indicating that cell division may give rise to daughter cells with distinct fates. Based on our observation that the RAF can be absent in the LN of a CLL patient with aggressive disease and expanded in the PB, we speculate that stimulation of CXCR4^hi^ cells in tissue gives rise to the RAF in the PB. Subsequent transition of cells from the RAF to the PF is likely dependent on the strength and nature of the initial stimulation with only highly stimulated cells progressing to S/G2/M phases and the remaining cells transitioning to more quiescent phenotypes. Additionally, we detected an expanded Ki67^hi^ PF in the LN and observed a similar phenomenon in the spleens of Eu-TCL1-tg mice that develop an aggressive CLL-like disease, suggesting that an expanded PF in tissue is indicative of increased cell turnover. Conversely, we observed expansions of the QF in indolent CLL and in the BM of Eu-TCL1-tg mice, where frequencies of proliferating cells are low.

Whilst high CD5 expression may reflect recent activation, our observation that CXCR4 levels rise gradually from G1 to G2/M, establish a strong link between CXCR4 expression and cell cycle progression in CLL cells. Cell cycle entry may influence CXCR4 dynamics as CXCR4 internalisation is detected 2hrs post BCR stimulation in CLL cells^26^, consistent with lymphocytes committing to the G1 stage 3hrs post a stimulatory event^27^. Subsequent CXCR4 upregulation on dividing cells likely requires the synthesis of new CXCR4 transcripts as the majority of intracellular CXCR4 levels are degraded ∼6hrs post stimulation^26^, in contrast to rapid recovery of CXCR4 expression in the absence of stimulation due to recycling of the receptor^28,16^. Fluctuations in CXCR4 expression on proliferating CLL cells may mirror the changes in CXCR4 levels on GC B cells where CXCR4 expression discriminates between proliferating and non-proliferating subsets in the dark (DZ) and light zones (LZ) respectively^29,30^ In support, and consistent with previous analysis^15^, we observe highest levels of AID in the RAF in CLL, enriched in G1 cells, analogous to restricted AID activity in early G1-phase B cells in the DZ^31^. Finally, GC B cells are most sensitive to BCR stimulation in G2/M phase^32^, while we find that the PF, expresses high levels of IgM, associated with more efficient BCR signalling^16^. Direct regulation of the cell cycle by CXCR4 has yet to be established in CLL although previous studies have implicated CXCR4 signalling in cell cycle regulation in solid tumours. CXCR4 silencing inhibits cell growth in human carcinoma cell lines^33^ and increased expression of CXCR4 in G2/M phases has been reported in breast cancer cell lines where proteomic analysis revealed crosstalk between CXCR4 and G2-M transition proteins such as PLK1 and *Cyclin B1*^34^. This suggests that rather than a surrogate marker for cell division, active CXCR4 signalling in CLL cells may be an important regulator of the mitotic process.

We detected no differences in the PF size between progressing and indolent cases whilst expanded QFs were observed in indolent cases suggesting the CXCR4^hi^CD5^hi^ phenotype of mitotic cells is highly transient with cells either rapidly returning to tissue for restimulation or downregulating CD5 to transition to the QF where they accumulate and die. Supportive of the first scenario, we detected highest levels of CCR7 and efficient migration towards CCL21, required for LN homing, on the PF. Alternatively, the accumulation of quiescent CXCR4^hi^ cells may obscure the PF, limiting its use as a prognostic indicator. Importantly, we consistently detected increased sizes of the PF in patients at relapse compared to matched untreated samples, indicating that tracking the outgrowth of the PF may serve as an early biomarker of clinical progression. Moreover, as genetic mutations likely arise during S phase, sequencing of cells in G2/M phase may reveal acquisition of new genomic alterations, as well as the genetics of the most proliferative clones.

Previous analysis alluded to a minor subfraction of PB CLL cells expressing high levels of CXCR4 and IgM, open to both migration to tissue and BCR signals^16^. Cells in the PF are strong candidates for this subpopulation where divided cells are uniquely poised to home to tissue for restimulation. Importantly, high expression of CCR7, CXCR5, CD49d and IgM suggest these cells can efficiently respond to a range of tissue chemokine gradients maximising cell-cell interactions^35,36^. Our findings may facilitate therapeutic strategies targeting the most dangerous CLL cell fractions and provide a strong rationale for the use of monoclonal antibodies for CCR7 and CXCR4 that are currently being trialled^37,38^.

## Acknowledgements

The authors thank all the patients who were willing to donate samples for this study and Stephen Devereux for critically reading the manuscript. This work was supported by grants from Leukaemia UK (2019/JGF/002), Blood Cancer UK (22006) (RH) and the Lymphoma Research Trust (LRT6147) (DF).

## References

1. Messmer BT, Messmer D, Allen SL, et al. In vivo measurements document the dynamic cellular kinetics of chronic lymphocytic leukemia B cells. J Clin Invest. 2005;115(3):755–764. doi:10.1172/JCI23409

2. Damle RN, Wasil T, Fais F, et al. Ig V Gene Mutation Status and CD38 Expression As Novel Prognostic Indicators in Chronic Lymphocytic Leukemia. Blood. 1999;94(6):1840–1847. doi:10.1182/BLOOD.V94.6.1840

3. Hamblin TJ, Davis Z, Gardiner A, Oscier DG, Stevenson FK. Unmutated Ig V(H) genes are associated with a more aggressive form of chronic lymphocytic leukemia. Blood. 1999;94(6):1848–1854. doi:10.1182/blood.v94.6.1848

4. Herishanu Y, Pérez-Galán P, Liu D, et al. The lymph node microenvironment promotes B-cell receptor signaling, NF-κB activation, and tumor proliferation in chronic lymphocytic leukemia. Blood. 2011;117(2):563–574. doi:10.1182/blood-2010-05-284984

5. Herndon TM, Chen SS, Saba NS, et al. Direct in vivo evidence for increased proliferation of CLL cells in lymph nodes compared to bone marrow and peripheral blood. Leukemia. 2017;31(6):1340–1347. doi:10.1038/leu.2017.11

6. Calissano C, Damle RN, Marsilio S, et al. Intraclonal complexity in chronic lymphocytic leukemia: Fractions enriched in recently born/divided and older/quiescent cells. Mol Med. 2011;17(11):1374–1382. doi:10.2119/molmed.2011.00360

7. Morande PE, Sivina M, Uriepero A, et al. Ibrutinib therapy downregulates AID enzyme and proliferative fractions in chronic lymphocytic leukemia. Blood. 2019;133(19):2056–2068. doi:10.1182/blood-2018-09-876292

8. Palacios F, Yan XJ, Ferrer G, et al. Musashi 2 influences chronic lymphocytic leukemia cell survival and growth making it a potential therapeutic target. Leuk 2021 354. 2021;35(4):1037–1052. doi:10.1038/s41375-020-01115-y

9. Haselager M V., Kielbassa K, ter Burg J, et al. Changes in Bcl-2 members after ibrutinib or venetoclax uncover functional hierarchy in determining resistance to venetoclax in CLL. Blood. 2020;136(25):2918–2926. doi:10.1182/blood.2019004326

10. Sun C, Chen YC, Zurita AM, et al. The immune microenvironment shapes transcriptional and genetic heterogeneity in chronic lymphocytic leukemia. Blood Adv. 2023;7(1):145. doi:10.1182/BLOODADVANCES.2021006941

11. Pasikowska M, Walsby E, Apollonio B, et al. Phenotype and immune function of lymph node and peripheral blood CLL cells are linked to transendothelial migration. Blood. 2016;128(4):563–573. doi:10.1182/BLOOD-2016-01-683128

12. Muñoz-Ruiz M, Pujol-Autonell I, Rhys H, et al. Tracking immunodynamics by identification of S-G2/M-phase T cells in human peripheral blood. J Autoimmun. 2020;112:102466. doi:10.1016/J.JAUT.2020.102466

13. Cuthill KM, Zhang Y, Pepper A, et al. Identification of proliferative and non-proliferative subpopulations of leukemic cells in CLL. Leukemia. 2022;36(9):2233. doi:10.1038/S41375-022-01656-4

14. D’Avola A, Drennan S, Tracy I, et al. Surface IgM expression and function are associated with clinical behavior, genetic abnormalities, and DNA methylation in CLL. Blood. 2016;128(6):816–826. doi:10.1182/BLOOD-2016-03-707786

15. Patten PEM, Chu CC, Albesiano E, et al. IGHV-unmutated and IGHV-mutated chronic lymphocytic leukemia cells produce activation-induced deaminase protein with a full range of biologic functions. Blood. 2012;120(24):4802–4811. doi:10.1182/blood-2012-08-449744

16. Coelho V, Krysov S, Steele A, et al. Identification in CLL of circulating intraclonal subgroups with varying B-cell receptor expression and function. Blood. 2013;122(15):2664–2672. doi:10.1182/blood-2013-02-485425

17. Cuesta-Mateos C, Terrón F, Herling M. CCR7 in Blood Cancers – Review of Its Pathophysiological Roles and the Potential as a Therapeutic Target. Front Oncol. 2021;11:736758. doi:10.3389/FONC.2021.736758/BIBTEX

18. Mazzarello AN, Fitch M, Cardillo M, et al. Characterization of the Intraclonal Complexity of Chronic Lymphocytic Leukemia B Cells: Potential Influences of B-Cell Receptor Crosstalk with Other Stimuli. Cancers (Basel). 2023;15(19):4706. doi:10.3390/CANCERS15194706/S1

19. Miller I, Min M, Yang C, et al. Ki67 is a Graded Rather than a Binary Marker of Proliferation versus Quiescence. Cell Rep. 2018;24(5):1105. doi:10.1016/J.CELREP.2018.06.110

20. Di Rosa F, Cossarizza A, Hayday AC. To Ki or Not to Ki: Re-Evaluating the Use and Potentials of Ki-67 for T Cell Analysis. Front Immunol. 2021;12:653974. doi:10.3389/FIMMU.2021.653974/BIBTEX

21. Sobecki M, Mrouj K, Camasses A, et al. The cell proliferation antigen Ki-67 organises heterochromatin. Elife. 2016;5(MARCH2016). doi:10.7554/ELIFE.13722

22. Sobecki M, Mrouj K, Colinge J, et al. Cell-cycle regulation accounts for variability in Ki-67 expression levels. Cancer Res. 2017;77(10):2722–2734. doi:10.1158/0008-5472.CAN-16-0707/652407/AM/CELL-CYCLE-REGULATION-ACCOUNTS-FOR-VARIABILITY-IN

23. Pauklin S, Vallier L. The Cell-Cycle State of Stem Cells Determines Cell Fate Propensity. Cell. 2013;155(1):135. doi:10.1016/J.CELL.2013.08.031

24. Rodríguez Varela MS, Mucci S, Videla Richardson GA, et al. Regulation of cyclin E1 expression in human pluripotent stem cells and derived neural progeny. Cell Cycle. 2018;17(14):1721. doi:10.1080/15384101.2018.1496740

25. Gu Y, Hu X, Liu X, et al. MCM6 indicates adverse tumor features and poor outcomes and promotes G1/S cell cycle progression in neuroblastoma. BMC Cancer. 2021;21(1):1–14. doi:10.1186/S12885-021-08344-Z/FIGURES/6

26. Vlad A, Deglesne PA, Letestu R, et al. Down-regulation of CXCR4 and CD62L in chronic lymphocytic leukemia cells is triggered by B-cell receptor ligation and associated with progressive disease. Cancer Res. 2009;69(16):6387–6395. doi:10.1158/0008-5472.CAN-08-4750/655038/P/DOWN-REGULATION-OF-CXCR4-AND-CD62L-IN-CHRONIC

27. Lea NC, Orr SJ, Stoeber K, et al. Commitment Point during G0→G1 That Controls Entry into the Cell Cycle. Mol Cell Biol. 2003;23(7):2351–2361. doi:10.1128/MCB.23.7.2351-2361.2003

28. Zhang Y, Foudi A, Geay J, et al. Intracellular Localization and Constitutive Endocytosis of CXCR4 in Human CD34+ Hematopoietic Progenitor Cells. Stem Cells. 2004;22(6):1015–1029. doi:10.1634/STEMCELLS.22-6-1015

29. Caron G, Le Gallou S, Lamy T, Tarte K, Fest T. CXCR4 Expression Functionally Discriminates Centroblasts versus Centrocytes within Human Germinal Center B Cells. J Immunol. 2009;182(12):7595–7602. doi:10.4049/JIMMUNOL.0804272

30. Weber TS. Cell cycle-associated CXCR4 expression in germinal center B cells and its implications on affinity maturation. Front Immunol. 2018;9(JUN). doi:10.3389/fimmu.2018.01313

31. Wang Q, Kieffer-Kwon KR, Oliveira TY, et al. The cell cycle restricts activation-induced cytidine deaminase activity to early G1. J Exp Med. 2017;214(1):49. doi:10.1084/JEM.20161649

32. Khalil AM, Cambier JC, Shlomchik MJ. B Cell Signal Transduction in Germinal Center B Cells is Short-Circuited by Increased Phosphatase Activity. Science. 2012;336(6085):1178. doi:10.1126/SCIENCE.1213368

33. Liang S, Peng X, Li X, et al. Silencing of CXCR4 sensitizes triple-negative breast cancer cells to cisplatin. Oncotarget. 2014;6(2):1020–1030. doi:10.18632/ONCOTARGET.2741

34. Liu S, Xie SM, Liu W, et al. Targeting CXCR4 abrogates resistance to trastuzumab by blocking cell cycle progression and synergizes with docetaxel in breast cancer treatment. Breast Cancer Res. 2023;25(1):1–20. doi:10.1186/S13058-023-01665-W/FIGURES/7

35. Bürkle A, Niedermeier M, Schmitt-Gräff A, Wierda WG, Keating MJ, Burger JA. Overexpression of the CXCR5 chemokine receptor, and its ligand, CXCL13 in B-cell chronic lymphocytic leukemia. Blood. 2007;110(9):3316–3325. doi:10.1182/BLOOD-2007-05-089409

36. Cuesta-Mateos C, Brown JR, Terrón F, Muñoz-Calleja C. Of Lymph Nodes and CLL Cells: Deciphering the Role of CCR7 in the Pathogenesis of CLL and Understanding Its Potential as Therapeutic Target. Front Immunol. 2021;12:927. doi:10.3389/fimmu.2021.662866

37. Kashyap MK, Amaya-Chanaga CI, Kumar D, et al. Targeting the CXCR4 pathway using a novel anti-CXCR4 IgG1 antibody (PF-06747143) in chronic lymphocytic leukemia. J Hematol Oncol. 2017;10(1). doi:10.1186/S13045-017-0435-X

38. Cuesta-Mateos C, Juárez-Sánchez R, Mateu-Albero T, et al. Targeting cancer homing into the lymph node with a novel anti-CCR7 therapeutic antibody: the paradigm of CLL. MAbs. 2021;13(1). doi:10.1080/19420862.2021.1917484

