## Supplementary figures for "Proliferating CLL cells express high levels of CXCR4 and CD5"

### Supplementary Figure 1.

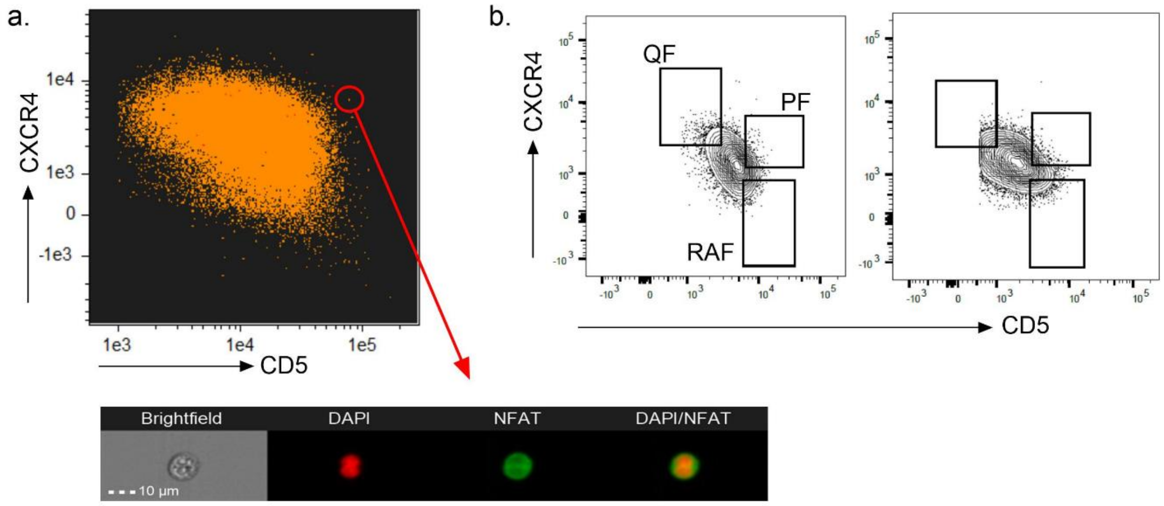


**Supplementary figure 1. Proliferating CLL cells can be observed in the CXCR4$^{hi}$CD5$^{hi}$ fraction.** (a) Image of a dividing CXCR4$^{hi}$CD5$^{hi}$ PB CLL cell in G2/M phase acquired using imaging flow microscopy. Red: DAPI (nucleus); Green: NFAT2 (cytoplasm). PB, peripheral blood. (b) Representative CXCR4/CD5 contour plots from 2 CLL patients demonstrating the gating strategy for the quiescent fraction (QF), proliferating fraction (PF) and recent activated fraction (RAF). Gates were set to capture 5% of the bulk population in each fraction.

### Supplementary Figure 2.

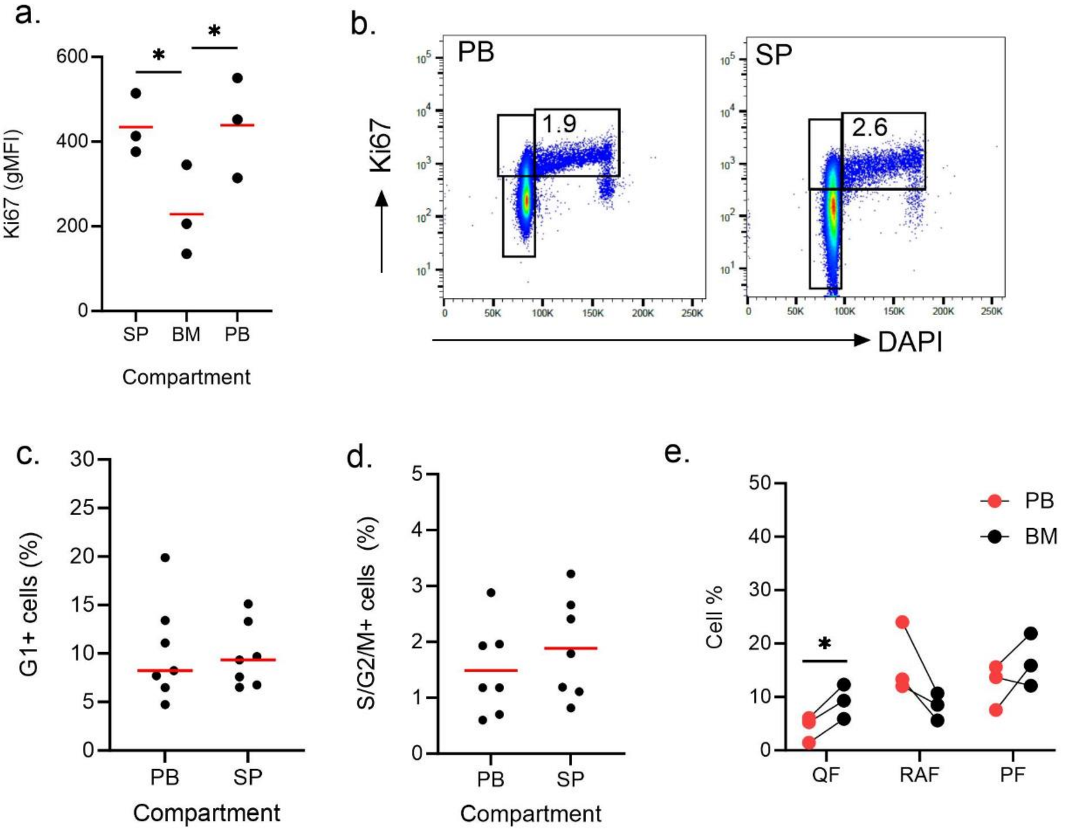


**Supplementary figure 2. The bone marrow does not sequester proliferating cells in the Eµ-TCL1 mouse model.** (a) Quantification of Ki67 levels in CD19+CD5+ TCL1 cells from the spleen (SP), bone marrow (BM) and peripheral blood (PB). Each data point represents a single mouse (n=3). (b) Representative scatter plots of Ki67 and DAPI staining of CD19+CD5+ cells in the PB and SP highlighting cell fractions in G0, G1 and S/G2/M phases. Quantification of the percentage of cells in (c) G1 and (d) S/G2/M between PB and SP compartments. (n=7) (e) Quantification of cell percentages in the three different fractions compared to matched PB fractions in the bone marrow (n=3). QF: Quiescent Fraction, RAF: Recent Activated Fraction, PF: Proliferating Fraction. Statistical significance of data were calculated using a RM one way ANOVA with Tukey's multiple comparisons or a paired parametric t-test. *p <0.05.

### Supplementary Figure 3.

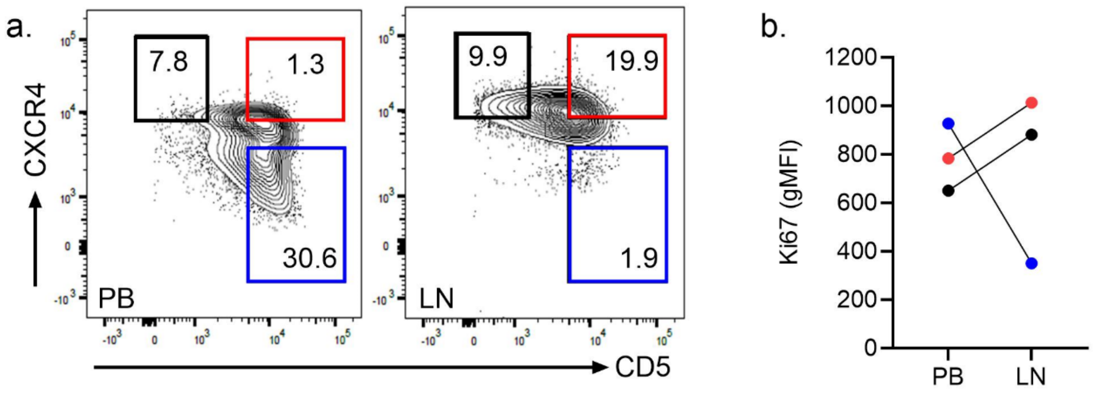


**Supplementary figure 3. An expanded CXCR4$^{hi}$CD5$^{hi}$ fraction is observed in the lymph nodes of a CLL patient with aggressive disease.** (a) CXCR4 and CD5 contour plots of LN and matched PB CLL cells from a de novo U-CLL patient with mutated TP53 and rapidly progressing disease. (b) Quantification of intracellular Ki67 with colours reflecting the different cell fractions gated in (a). gMFI, geometric mean fluorescence intensities.

### Supplementary Figure 4.

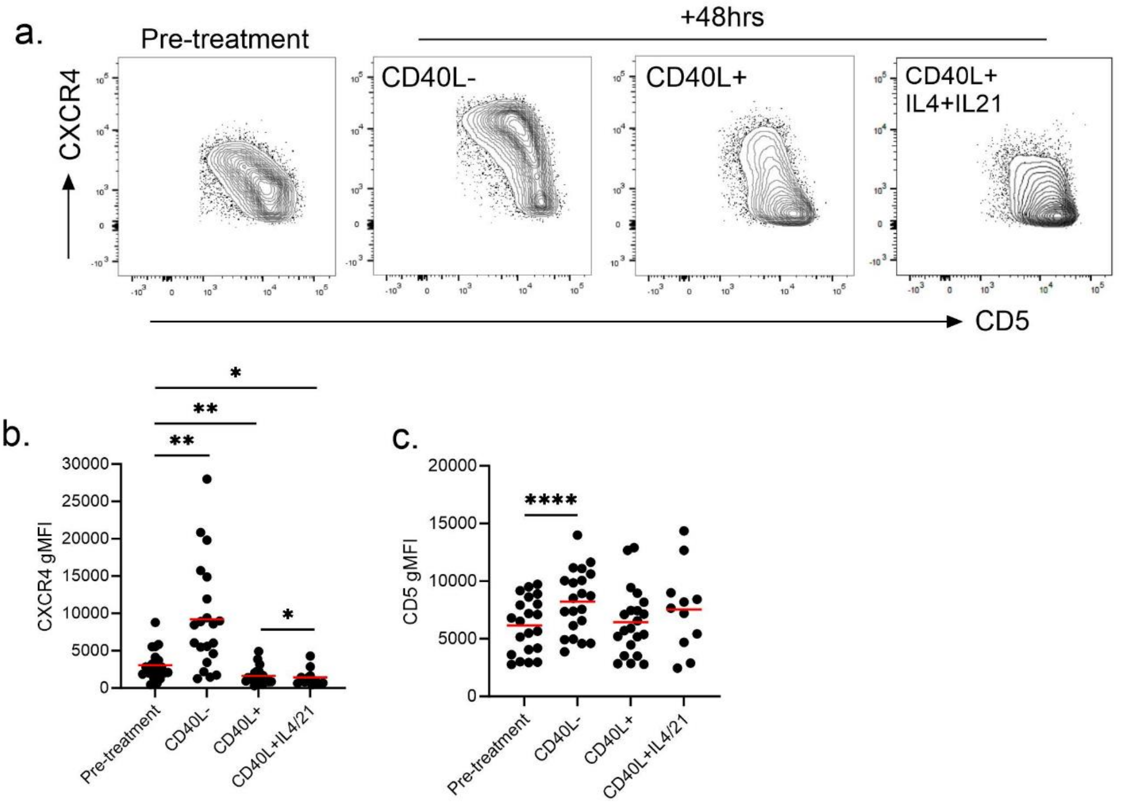


**Supplementary figure 4. CXCR4 expression is downregulated in response to activating signals.** (a) Human U-CLL PBMCs were seeded on CD40L-, CD40L+ fibroblasts alone or in the presence of interleukin (IL)-4 and IL-21 for 48 hrs and stained for CD19, CD5, and CXCR4. Representative contour CXCR4/CD5 plots from cells prior to treatment and after 48hrs in different conditions are shown. (b) Bulk CXCR4 and (c) CD5 levels were quantified on CLL cells both prior to treatment and after 48hrs stimulation (n=21). Data points represent individual patients. gMFI, geometric mean fluorescence intensities. Statistical significance of data were calculated using a RM one way ANOVA with Tukey's multiple comparisons. *p <0.05, **p <0.01, ****p <0.0001.

### Supplementary Figure 5.

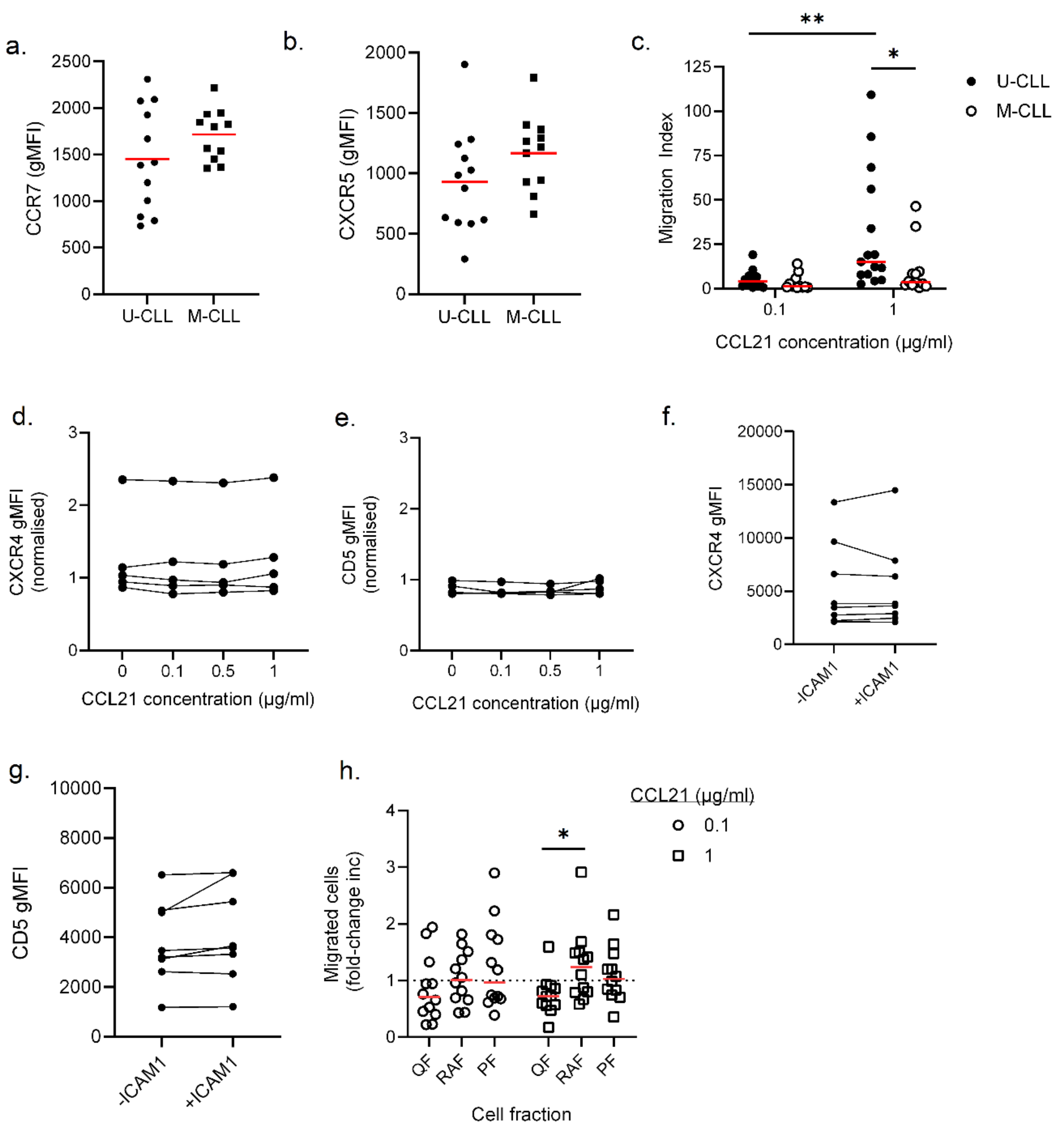

**Supplementary figure 5. CXCR4 and CD5 levels do not fluctuate in response to CCR7 signalling.** Bulk expression levels of (a) CCR7 and (b) CXCR5 were quantified on U- and M-CLL cells. (c) PBMCs from both U- and M-CLL patients were placed in transwell migration chambers and incubated in the absence or presence of increasing concentrations of CCL21. After 2hrs, migrated cells were stained with CD19 and CD5 and numbers quantified by flow cytometry. Migration index = number of migrated cells with chemokine/number of migrated cells in the absence of chemokine. (d) PBMCs were incubated with increasing concentrations of CCL21 for 2hrs at 37°C and CXCR4 and (e) CD5 levels quantified by flow cytometry (n=5). (f) PBMCs were seeded on ICAM-1 coated well plates for 2hrs at 37°C and CXCR4 and (g) CD5 levels quantified by flow cytometry (n=8) (h) PBMCs from M-CLL patients were placed in transwell migration chambers and incubated for 2hrs at 37°C in the absence or presence of increasing concentrations of CCL21. After 2hrs migrated cells were harvested from the bottom chamber and stained with antibodies against CD19, CD5, CXCR4 and CD5. CXCR4 and CD5 gates were drawn on time matched controls and extrapolated on migrated fractions to quantify changing fraction sizes. Quantification of the fold-change increase in fraction size in M-CLL patients of migrated cells is shown at both lower (0.1µg/ml) and higher (1µg/ml) CCL21 concentrations (n=12). Data points represent individual patients. QF: Quiescent Fraction, RAF: Recent Activated Fraction, PF: Proliferating Fraction. Statistical significance of data were calculated using a RM one way ANOVA with Tukey's multiple comparisons or an unpaired parametric t-test. *p <0.05, **p <0.01.
